# Reduced Myocardial Serine Synthesis Impairs Functional, Metabolic, and Redox Adaptations to Cardiac Stress

**DOI:** 10.64898/2026.05.29.728910

**Authors:** Malihe Rezaee, Mohammad Keykhaei, Navid Koleini, Tegbir Panesar, Simiao Li, Nalini Salvekar, David J Polhemus, Chengchen Hu, Mariam Meddeb, Liang Zhao, Kavita Sharma, Christopher Petucci, Nathaniel Snyder, Junichi Sadoshima, David A. Kass

## Abstract

**Background:** Impaired myocardial metabolism is a defining feature of heart failure, but many defective pathways and mechanisms remain to be identified. Prior studies find phosphoglycerate kinase and its synthesized product 3-phospho-glycerate required for the serine synthetic pathway (SSP) are reduced in human HFpEF myocardium. As serine is also provided exogenously, the impact of SSP reduction is uncertain. Here, we tested if and how SSP decline coupled to phosphoglycerate dehydrogenase (PHGDH) impacts cardiomyocyte (CM) and whole heart metabolic remodeling and stress responses.

**Methods:** Studies were performed in isolated CMs and mice with CM-selective knock-down of PHGDH. Using pharmacological inhibition or genetic silencing of PHGDH, we tested their impact on CM one-carbon metabolism pathways, cell hypertrophic responses, mitochondrial respiration, and *in vivo* functional, structural, and metabolic adaptations to pressure-overload stress.

**Results:** In CMs, PHGDH inhibition caused dose-dependent serine depletion linearly coupled with cytotoxicity, accompanied by NAD/NADH and GSH/GSSG imbalance, reduced ATP, and disruption of one-carbon and nucleotide metabolites. Stable-isotope tracing revealed distinct metabolic fates of glucose-derived (SSP) versus exogenous serine. Exogenous serine did not rescue PHGDH-deficient CMs, whereas combined ribose and an anti-oxidant (DTT) attenuated injury and reduced nucleotide pools. PHGDH suppression reduced amino acid abundance, impaired nascent protein synthesis, and blunted endothelin-1–induced hypertrophic and mitochondrial respiration. *In vivo*, cardiomyocyte-specific PHGDH heterozygous mice (PHGDH^+/−^) had no basal phenotype, but amplified chamber dilation, dysfunction, fibrosis, and mortality 4 weeks after transverse aortic constriction (TAC). Corresponding increases in amino acids, one-carbon metabolites, nucleotides, and TCA-cycle intermediates in wild-type TAC hearts were significantly blunted in PHGDH^+/−^ hearts.

**Conclusions:** Cardiomyocyte SSP is a critical regulator of redox balance, one-carbon metabolism, purine synthesis, amino acid homeostasis, and growth-related pathways required for cardiac adaptation to pressure overload. It is non-redundant with exogenous serine by providing distinct influences on key metabolic pathways and is a potential therapeutic target.

## Introduction

Heart failure (HF) is a leading cause of morbidity and mortality and increasing in prevalence worldwide^1^. Over the past several decades, the characteristics of HF have evolved, with an increasing proportion of patients presenting with chronic obesity and metabolic disease. The shift particularly affects those with HF and a preserved ejection fraction (HFpEF)^2^, but it is also common in HF with reduced EF (HFrEF)^3^. Recent success in treating both forms of HF with drugs developed for diabetes, such as sodium-glucose transporter inhibitors^4^ and glucagon-like peptide receptor agonists^5^, adds support for the importance of metabolic abnormalities. This includes impaired myocardial fuel use and flexibility, carbohydrate and fatty acid metabolism, and inadequate generation of nucleotides needed for mitochondrial respiration and energy production^6^. Finding root causes for these defects to identify better targeted therapies remains a major goal.

We first reported that glycolysis intermediates are much reduced in human HFpEF myocardium^7^, and chief among these was 3-phosphoglycerate (3PG), the input substrate for *de novo* serine biosynthesis (SSP). In another study, we found myocardial serine is also reduced ^8^, while in the human HFrEF heart, changes in both 3PG and serine are reportedly less^8,9^. Serine is a non-essential amino acid that also enters cells via transporters or is generated by the SSP^10^. The latter converts 3PG into serine in three steps, the first and rate-limiting enzyme being phosphoglycerate dehydrogenase (PHGDH), followed by phosphoserine aminotransferase (PSAT1) and phosphoserine phosphatase (PSPH) ^11^. Beyond its role in protein composition, serine serves as a major metabolic node linking glycolysis to nucleotide and amino acid biosynthesis, lipid metabolism, and cellular antioxidant defenses ^12,13^. Through its involvement in one-carbon metabolism^14^ (folate, methionine, and transsulfuration pathways), serine provides antioxidant and nucleotide pools for gene transcription, cell growth, redox balance, purine and thymidine synthesis, methylation by S-adenosyl methionine (SAM), and glutathione generation ^15–17^. These are particularly important in stressed conditions.

While mostly studied in cancer biology^18^, emerging evidence shows serine and the SSP are relevant to heart disease. SSP activation rescues contractile defects of induced pluripotent stem cell (iPSC)-derived cardiomyocytes derived from patients with genetic HFrEF ^19^, and *Phgdh* overexpression delivered by adeno-associated virus *in vivo* reverses pathological remodeling in a genetic HFrEF mouse model^20^. Reciprocally, SSP inhibition depresses cardiomyocyte proliferation and triggers apoptosis after myocardial infarction^21^, and in iPSCs, it favors cardiomyocyte specification coupled to lower mitochondrial respiration and higher oxidant stress^22^. Still, much remains unknown about the relative role of SSP in cardiomyocytes and cardiac pathophysiology.

Accordingly, the present study was designed to determine the metabolic influences of cardiomyocyte SSP, its distinction from the exogenous serine pathway, and how its reduction impacts CM and whole heart stress adaptations. We identify PHGDH reduction in human HF, particularly HFpEF. The findings reveal that PHGDH-mediated serine biosynthesis is a major regulator of metabolic, structural, and functional cardiomyocyte biology, and its suppression contributes to heart failure and can be a therapeutic target.

## Materials and Methods

### Human Myocardial Tissue

Human endomyocardial biopsies (2-3 mg) were obtained from the right mid-ventricular septum of patients enrolled at the Johns Hopkins Heart Failure with Preserved Ejection Fraction (HFpEF) Clinic under Institutional Review Board-approved protocols. Written informed consent was obtained from all participants. The HFpEF cohort was well phenotyped, including right heart catheterization and echocardiography and met consensus diagnostic criteria^7,8^. Biopsies were rapidly frozen in liquid nitrogen and maintained at -160°C until analyzed. Non-failing (NF) control myocardium was from organ donors without known cardiac disease whose hearts were not used for transplantation. Heart excision was done using the same surgical cardioplegic arrest techniques for HFrEF transplant recipients. All tissue samples were fully de-identified before analysis.

### Bulk RNA Sequencing and Metabolomic Analysis of human heart tissues

Bulk RNA sequencing methods and analyses of HFpEF, HFrEF, and NF control human myocardium have been previously reported ^23^, and targeted gene expression analysis in the current study was derived from this prior work. Detailed methods for myocardial and plasma metabolites in the same three patient groups have also been previously reported ^8^. Additional *de novo* metabolic analyses are described below.

### Neonatal Rat Ventricular Myocyte (NRVM)

Primary cardiomyocyte experiments were conducted using neonatal rat ventricular myocytes (NRVMs) isolated from 1–2-day-old Sprague–Dawley pups by established enzymatic dissociation methods ^24^. Methods for siRNA modification of NRVMs are provided in Supplemental Methods.

### Adult Mouse Cardiomyocytes

Adult mouse ventricular cardiomyocytes were isolated from 8–12-week-old mice using a Langendorf-free retrograde coronary perfusion approach as described ^25^.

### Immunoblots for protein expression analysis

Details of protein expression analysis by Western blot, including specific antibodies and titers, are provided in the supplemental methods.

### RNA Extraction and Quantitative PCR

Expression of mRNA analysis followed standard protocols detailed in Supplemental Methods.

### Myocyte pharmacological treatments

For pharmacological inhibition of PHGDH, NCT-503 (Sigma-Aldrich; CAS 1916571-90-8) was administered at 1, 2, 3, or 10 µmol/L for 24 h for LDH release and serine quantification assays. For all other inhibitor studies, NCT-503 was used at 10 µmol/L for 24 h. For rescue experiments, cells were treated for 24 h with sodium pyruvate (Santa Cruz Biotechnology; CAS 113-24-6; 1 mmol/L), L-serine (Sigma-Aldrich, S4311; 0.4 mmol/L), glycine (Sigma-Aldrich, G8790; 0.4 mmol/L), MEM amino acids solution (50×; Gibco, 11130051; final 2×), β-nicotinamide mononucleotide (NMN; Cayman Chemical, 33732; 1 mmol/L), D-(−)-ribose (Sigma-Aldrich, R9629; 2 mmol/L), dithiothreitol (DTT; Thermo Fisher Scientific, D1532; 0.3 mmol/L), or EmbryoMax nucleosides (100×; Sigma-Aldrich, ES-008-D; final 1×). Media pH was adjusted to 7.4 after the addition of each supplement. For hypertrophic stimulation studies, NRVMs were exposed to endothelin-1(ET-1) (Sigma-Aldrich, E7764) at 100 nmol/L for 24 h.

### Nascent protein synthesis assay

Newly synthesized protein production was quantified using a puromycin-labeling assay. Cells were exposed to puromycin (Sigma, #P8833) at 10 µmol/L for 30 min to permit incorporation into elongating peptide chains. Cells were then washed with PBS, lysed, and the resulting protein samples subjected to immunoblot analysis.

### Metabolite Extraction and Mass Spectrometry Analysis

Details of the extraction procedure, and targeted mass spec analysis of amino acids, nucleotides, 1-carbon metabolites, glycolysis, and TCA intermediates were quantified using targeted liquid chromatography–tandem mass spectrometry (LC–MS/MS). Details in Supplemental Methods. The ^13^C₆-glucose and ^13^C₃-serine tracing analysis is described in Supplemental Methods.

### Mitochondrial oxidative respiration analysis

Cellular bioenergetics were assessed using the Seahorse XF Extracellular Flux Analyzer (Agilent Technologies) according to the manufacturer’s instructions (Supplemental Methods).

### Mouse model for heterozygote PHGDH knockdown using Myh6-Cre

PHGDH^fl/fl^ mouse embryos were obtained from RIKEN Bioresource Research Center (BRC No. RBRC02574) and rederived at Johns Hopkins Mouse Model Core. They were then crossed with *Myh6*-*Cre* mice (C57BL/6 background, Jax #011038) to generate a cardiomyocyte-specific heterozygote PHGDH knockout mouse (PHGDH^+/-^ and PHGDH^fl/fl^ littermate controls. Mice were housed in a temperature-controlled environment (21 to 23°C) with a 12-hour light/12-hour dark cycle and provided free access to water and a standard diet. For heart sample collection, mice were euthanized by cervical dislocation. In a separate set of 4-week TAC experiments, PHGDH^+/-^ were crossed with *Myh6*-MerCreMer^26^, and gene knockdown was induced by tamoxifen (Inotiv, TD.130855) administered for 7 days beginning 3 weeks after TAC. Tamoxifen-treated *Phgdh^flox/+^* mice lacking Cre recombinase were controls. Animal procedures were done in accordance with NIH guidelines and protocols approved by the Animal Care and Use Committee of Johns Hopkins University and Rutgers University. Animal experiments were performed in accordance with protocols approved by the Johns Hopkins University and Rutgers University Animal Care and Use Committee.

### Pressure Overload Surgery (trans-aortic constriction, TAC)

Two-three month old mice were randomly assigned to either TAC or sham surgery groups performed as previously described ^27^ (details in Supplemental Methods).

### Echocardiography

Transthoracic M-mode echocardiography was performed in conscious mice using a Vevo 2100 imaging system (VisualSonics) equipped with an 18–38 MHz transducer, as described ^28^, and in another cohort, by a Vevo 3100 system equipped with a 30-MHz linear array transducer (FUJIFILM VisualSonics) ^29^ under isofluorane sedation (3% induction, 1-2% maintenance).

### Histology and Fibrosis Analysis

Heart specimens were fixed overnight at 4°C in 10% formalin prepared in PBS, then transferred to 70% ethanol for dehydration and processed for paraffin embedding. Transverse left ventricular sections (4 µm) were cut and mounted onto positively charged glass slides. Interstitial and replacement fibrosis were evaluated using Masson’s Trichrome staining (Richard-Allan Scientific, #87020) according to the manufacturer’s protocol. Whole-heart sections were digitally imaged, and the fibrotic area was quantified using ImageJ (Fiji) software.

### Statistics

Statistical analyses were performed using GraphPad Prism version 11 (GraphPad Software). Sample size, statistical test and associated P-value, and multiple comparisons test used for each figure panel is provided in Supplemental Table. P-values for significant comparisons are provided in the individual figure. Individual biological replicates are shown in the figures unless otherwise indicated. Data are shown as mean±SD unless stated in the legend. Normality of the distributions was assessed by Kolmogorov–Smirnov test, and for 2-group comparisons, an unpaired 2-tailed Student’s *t* test or Mann–Whitney test was used. For >2 groups, 1-way or 2-way ANOVA, with Welch’s ANOVA used if unequal variances were identified by the Brown–Forsythe test. Nonparametric comparisons across multiple groups were performed using the Kruskal–Wallis test with Benjamini-Hochberg multiple comparison correction. Survival was analyzed by the Kaplan–Meier method, and differences between groups were assessed using the log-rank test. Exact *P* values are provided in the figures.

## Results

### Depressed serine, PHGDH, and one-Carbon metabolism in human heart failure

Figure 1A summarizes the exogenous and SSP serine pathways and related downstream metabolic pathways examined in this study. Serine levels in human myocardial tissue from NF control, HFpEF, and HFrEF are shown in Fig. 1B. Both HF conditions had reduced levels, HFpEF being significantly below HFrEF. However, plasma serine levels were similarly elevated in both HFpEF and HFrEF, indicating a dissociation between circulating and myocardial serine pools in HF.

**Figure 1:**
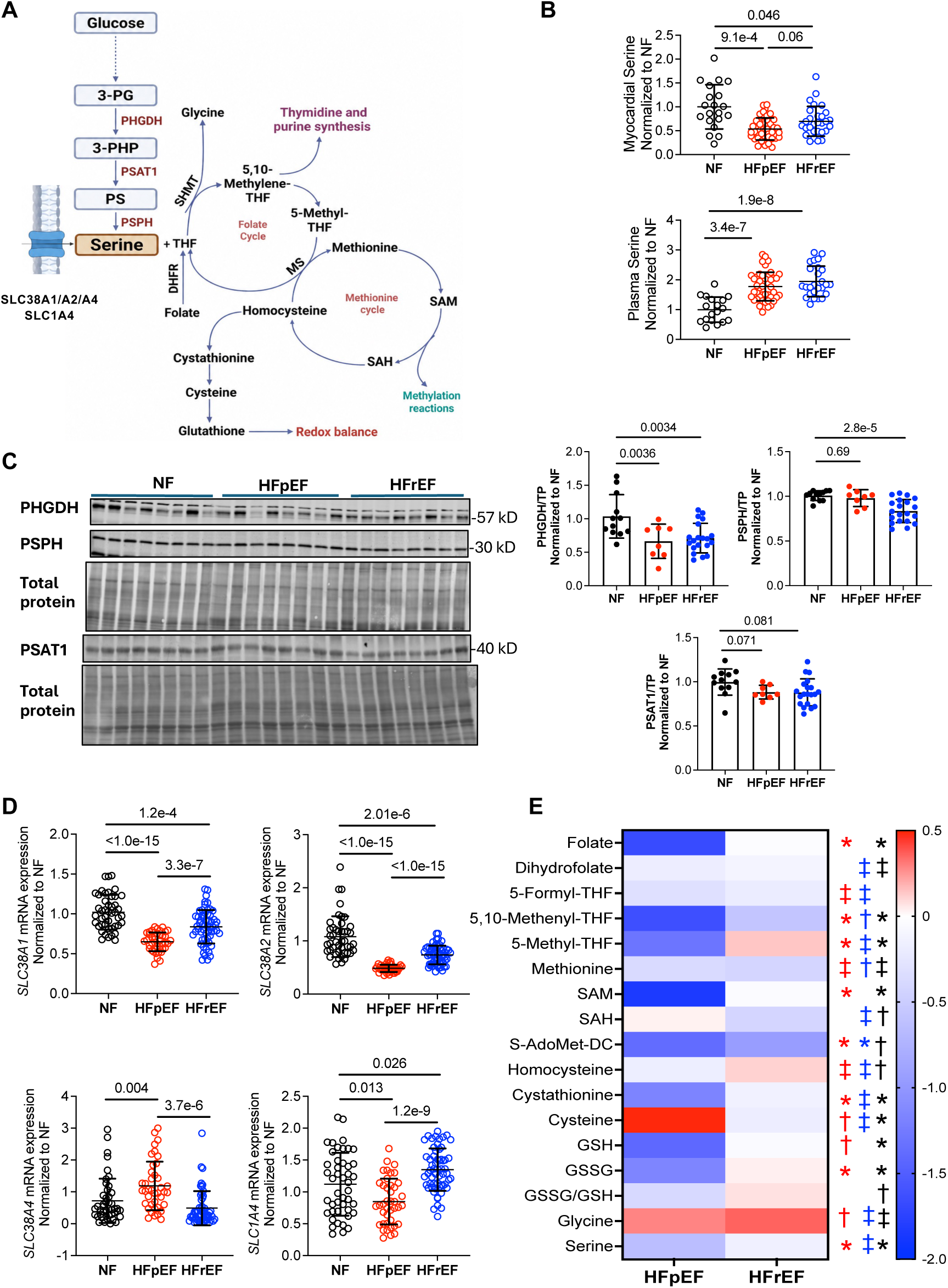
Myocardial serine and one-carbon metabolism are disrupted in human HFpEF. **A)** Schematic of glycolysis-linked endogenous serine biosynthesis and its coupling to one-carbon metabolism. 3-PG: Glucose-derived 3-phosphoglycerate 3-PHP: 3-phosphohydroxypyruvate, PHGDH: phosphoglycerate dehydrogenase, PS: phosphoserine, PSAT1: by phosphoserine aminotransferase 1, PSPH: phosphoserine phosphatase. **B)** Myocardial and plasma serine abundance in non-failing controls (NF), heart failure with preserved or reduced ejection fraction (HFpEF, HFrEF). Data are normalized to NF. **C)** Immunoblot of myocardial expression of PHGDH, PSPH, PSAT1, and total protein in three groups and summary densitometry, data normalized to total protein. **D)** Myocardial mRNA abundance of CM-expressed serine transporters in three patient groups. **E)** Heat map for levels of one-carbon-related metabolites for HFpEF and HFrEF normalized to NF control. Comparisons for HFpEF versus NF (red), HFrEF versus NF (blue), HFpEF versus HFrEF (black) are shown. P<1×10⁻⁶; †P<1×10⁻²; ‡P<0.09.

We next determined whether reduced myocardial serine was coupled with lower protein expression of SSP enzymes. PHGDH was significantly lower (P≤0.005) in both HFrEF and HFpEF, whereas PSPH was mildly reduced and only in HFrEF, and PSAT1 was borderline lower (P≤0.08) in both HF groups (Figure 1C). Corresponding mRNA expression for *Phgdh* and *Psph* was higher both HF groups versus NF (*Psat1* unchanged; Supplementary Figure 1). Gene expression for serine plasma membrane transporters most expressed in cardiomyocytes, SLC38A1, SLC38A2, was reduced in both HF conditions, but more so in HFpEF. Two other transporters, SLC38A4 and SLC1A4, either slightly increased or were unchanged (Figure 1D). Thus, both depressed SSP and serine transporters are particularly present in human HFpEF.

Targeted metabolic profiling for folate and methionine cycle and transsulfuration metabolites is presented in Figure 1E. Folate, dihydrofolate, and 5,10-methenyl-THF were most markedly reduced in HFpEF, accompanied by less methionine, SAM, and decarboxy-S-adenosylmethionine (dcSAM, the latter critical for polyamine synthesis) (red symbols). Several of these changes were shared by HFrEF (blue symbols), but the magnitude was generally less (black). There were also changes in transsulfuration and redox pathways; cystathionine, GSH, and GSSG all declined in HFpEF. Myocardial glycine was higher in both forms of HF, while serine was again found lower in HFpEF.

### PHGDH suppression induces cardiomyocyte injury coupled to serine reduction

Since PHGDH was most reduced in human HF myocardium, we first explored loss-of-function effects in cardiomyocytes with or without exogenous serine and glycine (S/G). Inhibiting PHGDH for 24 hours with NCT-503 in neonatal rat ventricular myocytes (NRVMs) concomitantly supplied exogenous S/G yielded a dose-dependent decline in intracellular serine that strongly and linearly correlated with LDH release, a measure of cell stress (Figure 2A). Serine decline did not induce cell death (Supplemental Figure 2A). This result indicates that despite external sources, sustained suppression of the SSP was sufficient to compromise serine levels in cardiomyocytes. The data also support the use of LDH as a surrogate biomarker for serine depletion in this setting. Time-course data using the NCT-503 dose that lowered serine by 75% showed no change in LDH until 24 hours (Figure 2B), while cells in S/G-free media had a similar rise by 4 hours (Figure 2B). Similar results were obtained in adult cardiomyocytes (Supplementary Figure 2B and 2C). PHGDH protein abundance was also diminished by siRNA (over 48 hours, Figure 2C). This induced a similar rise in LDH release as seen with NCT-503 in myocytes cultured in complete media, and the response nearly doubled in cells lacking exogenous S/G (Supplemental Figure 2D), again without evidence of cell death (Supplemental Figure 2E). As the substrate of PHGDH, 3PG, was also reduced in HFpEF myocardium, we tested the impact of genetic knockdown of phosphoglycerate kinase, the enzyme responsible for generating 3PG, by siRNA (Supplemental Figure 3A). This increased LDH much like with PHGDH-KD, and the combination was additive (Figure 2D). Elevating glucose, however, did not overcome PHGDH-KD (Figure 2E), indicating that selective glycolytic control was required.

**Figure 2:**
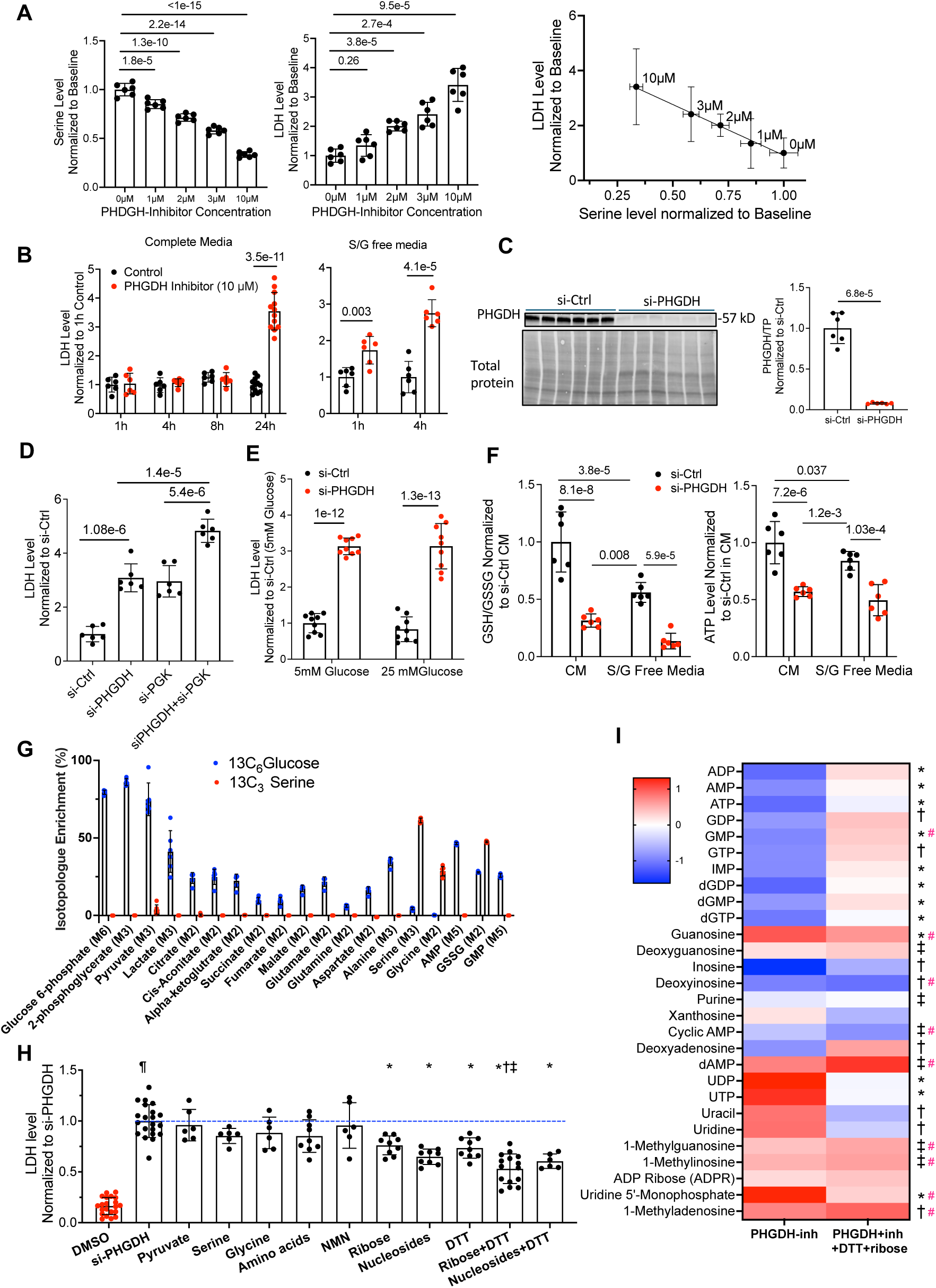
Effects of PHGDH suppression on serine, cell viability, redox, and nucleotide metabolism. **A)** *Left:* Serine level and LDH release following 24-hour treatment with increasing concentrations of PHGDH inhibitor NCT-503 in NRVMs. Data are normalized to respective control values. *Right:* Serine vs LDH plot reveals strong linear dose-dependence. **B)** LDH release from NRVMs treated with 10 µM NCT-503 in complete medium (CM) or in serine/glycine (S/G)-free medium for the indicated time durations. **C)** Immunoblot showing PHGDH expression and total protein in NRVMs treated with control siRNA (si-Ctrl) or PHGDH siRNA (si-PHGDH), with summary densitometry normalized to si-Ctrl. **D)** LDH release from NRVMs exposed to scrambled si-control, si-PHGDH, phosphoglycerate kinase knockdown (si-PGK), or combined PHGDH and PGK KD. **E)** LDH release in si-Ctrl or si-PHGDH-treated NRVMs cultured in 5 mmol/L or 25 mmol/L glucose. **F)** GSH/GSSG ratio and ATP abundance in si-Ctrl and si-PHGDH NRVMs cultured in complete medium (CM) or serine/glycine-free medium (S/G). Data normalized to si-Ctrl in CM control. **G)** Carbon labeling of metabolites after 24 hours of incubation with ^13^C_6_-glucose or ^13^C_3_-serine in NRVMs. Peaks with the anticipated number of labeled carbons are shown. **H)** LDH release in si-PHGDH-treated NRVMs with concomitant addition of either pyruvate, serine, glycine, amino acids, β-nicotinamide mononucleotide (NMN), ribose, nucleosides, dithiothreitol (DTT), ribose+DTT, and nucleosides+DTT; data normalized to si-PHGDH. * *P*<0.05 between condition and si-PHGDH; ¶ versus DMSO control, † versus ribose, and ‡ versus DTT. **I)** Heat map of nucleotide and nucleoside metabolites in NRVMs treated with NCT-503 versus NCT-503+ DTT and ribose, each normalized to untreated controls. Black symbols indicate significant differences between control and NCT503-treated cells: *P*<1×10⁻⁴; †*P*<5×10⁻³; ‡*P*<0.05. Red # indicates significant differences between NCT503-treated cells and NCT503-treated cells supplemented with DTT+ribose; #*P*<0.05.

### PHGDH reduction disrupts myocyte redox, nucleotide levels, and metabolic networks

Given the central role of serine metabolism in redox homeostasis and nucleotide synthesis^10^, downstream metabolic consequences of PHGDH suppression were examined in CMs. PHGDH-KD reduced the NAD^+^/NADH and oxidized/reduced glutathione (GSH/GSSG) ratios, and total ATP (Figure 2F, Supplemental Figure 3B). Intriguingly, reduction of the latter exceeded that after exogenous S/G removal (Figure 2F), suggesting different fates of the serine provided by the SSP vs exogenous sources. To explore this further, CMs were incubated with ^13^C_6_-glucose or ^13^C_3_-serine, and labeled-carbon tracing performed by mass spectrometry (Figure 2G). Only ^13^C_6_-glucose labeled glycolytic and TCA cycle intermediates, as well as selected amino acids. The latter included glutamate, a component of GSSG. Both purines, AMP and GMP, were labeled primarily from glucose. By contrast, ^13^C_3_-serine labeling was enriched in serine and glycine, less dominantly in GSSG, and not at all in TCA or glycolytic intermediates.

The impact of PHGDH-KD on 1-CM metabolites was examined by mass spectrometry (Supplemental Figure 3C), which revealed decreased methionine-cycle and folate-cycle metabolites, as well as GSH, with an increased GSSG/GSH ratio consistent with oxidant stress. To determine mechanisms most related to cytotoxicity from PHGDH-KD, a rescue experiment was performed (Figure 2H). Medium supplementation with pyruvate, serine, glycine, amino acids, or NMN (an NAD precursor) did not significantly lower LDH in PHGDH-KO CMs. However, ribose, nucleosides, and the reducing agent DTT each partially attenuated LDH, and combined ribose+DTT or nucleosides+DTT yielded the best rescue. These data highlight impaired nucleotide metabolism and redox buffering as major detrimental consequences of PHGDH deficiency in CMs. That exogenous serine failed to rescue the phenotype indicates that CMs preferentially rely on SSP-derived serine for these pathways. Based on this, we performed targeted nucleotide profiling in CMs exposed to PHGDH inhibitor ± ribose+DTT (Figure 2I). PHGDH inhibition broadly reduced purines and increased pyrimidines and adding ribose+DTT reversed most of these changes back towards control.

### PHGDH suppression broadly depresses amino acid levels and protein synthesis

The ^13^C-labeling study showed carbons from glucose and serine differentially resided in selected amino acids. We therefore explored the impact of PHGDH-KD on all amino acid levels by targeted mass spectrometry in CMs cultured in complete media ±siPHGDH, or in S/G depleted media, each for 48 hours (Figure 3A). This confirmed a greater decline in both serine and glycine with exogenous S/G depletion and the opposite pattern for alanine, each as predicted by the ^13^C-labeling study. Interestingly, both endogenous (SSP) and exogenous serine depletion led to reduced essential and non-essential amino acids (AAs), the latter suggesting either that their import was less important to intracellular levels in this setting, or that serine depletion impaired their importation.

**Figure 3:**
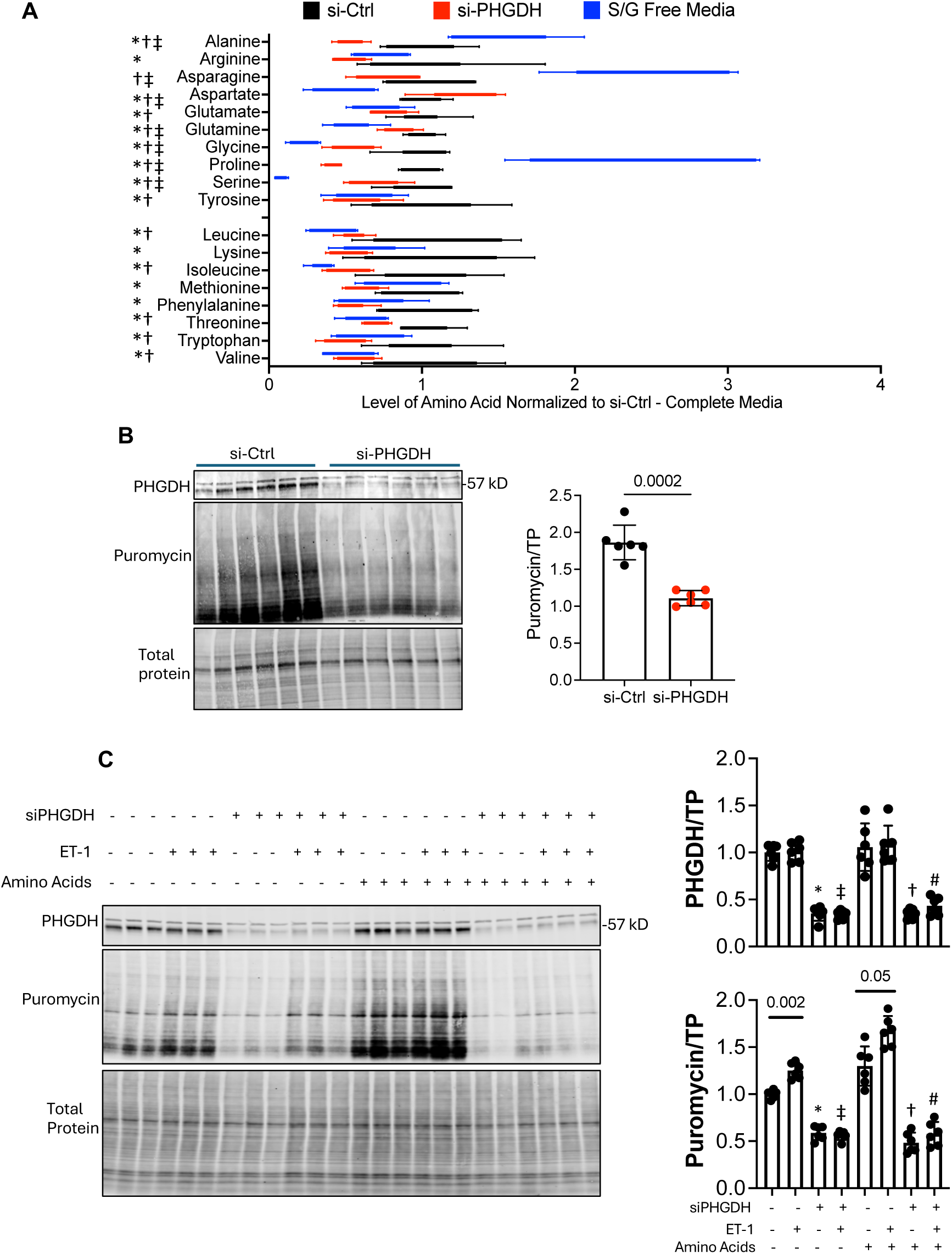
PHGDH suppression alters amino acid abundance and impairs nascent protein synthesis in NRVMs. **A)** Amino acid abundance in NRVMs treated with control siRNA (si-Ctrl), PHGDH siRNA (si-PHGDH) in complete media, or cultured in serine/glycine (S/G)-free media for 48 hours. Data normalized to si-Ctrl in complete media. Symbols indicate significant differences (*P*<0.05) between groups: *si-Ctrl versus si-PHGDH, †si-Ctrl versus S/G-free media, and ‡si-PHGDH versus S/G-free media. **B)** Immunoblot showing PHGDH expression, protein puromycin incorporation, and total protein in NRVMs treated with si-Ctrl or si-PHGDH for 48 hours. Summary densitometry for puromycin normalized to total protein on right. **C)** Immunoblot showing PHGDH, puromycin incorporation, and total protein in NRVMs treated with si-Ctrl or si-PHGDH, with or without endothelin-1 (ET-1) co-stimulation and with or without amino acid (AA) supplementation. Summary densitometry shown on the right (normalized to total protein). *P*<0.05 between * siPHGDH vs control, ‡ siPHGDH with ET-1 vs control with ET-1, † siPHGDH with AA vs control with AA, # siPHGDH with AA and ET-1 vs control with AA and ET-1.

The broad diminution in AAs from PHGDH-KD suggested protein synthesis may also become impaired. Nascent protein synthesis was measured by puromycin labeling assay, and it declined after PHGDH-KD (Figure 3B). This persisted in myocytes in which protein synthesis was stimulated by the pro-hypertrophic hormone endothelin-1 (ET-1) and was not restored despite supplementing exogenous AAs (Figure 3C). Thus, PHGDH suppression has a prominent and broad impact on amino acid abundance that impacts anabolic protein synthesis independent of exogenous amino acids.

### Inhibiting PHGDH blunts myocyte hypertrophic and metabolic response to ET-1

The preceding findings suggested that reduced PHGDH may impair physiological and molecular hypertrophic responses in CMs. This was tested by stimulating NRVMs with ET-1 that increased *Phgdh* expression and induced myocyte hypertrophy in control CMs, but this response was suppressed by PHGDH-KD (Figure 4A, 4B). Expression of mRNA markers of pathological hypertrophy: *Nppa*, *Nppb*, *Myh7*, and *Acta1* increased with ET-1 in controls, and each was attenuated by PHGDH-KD (Figure 4C). This was not due to greater cytotoxicity, as LDH augmentation with in PHGDH-KD was identical despite ET-1 stimulation (Supplementary Figure 3D).

**Figure 4:**
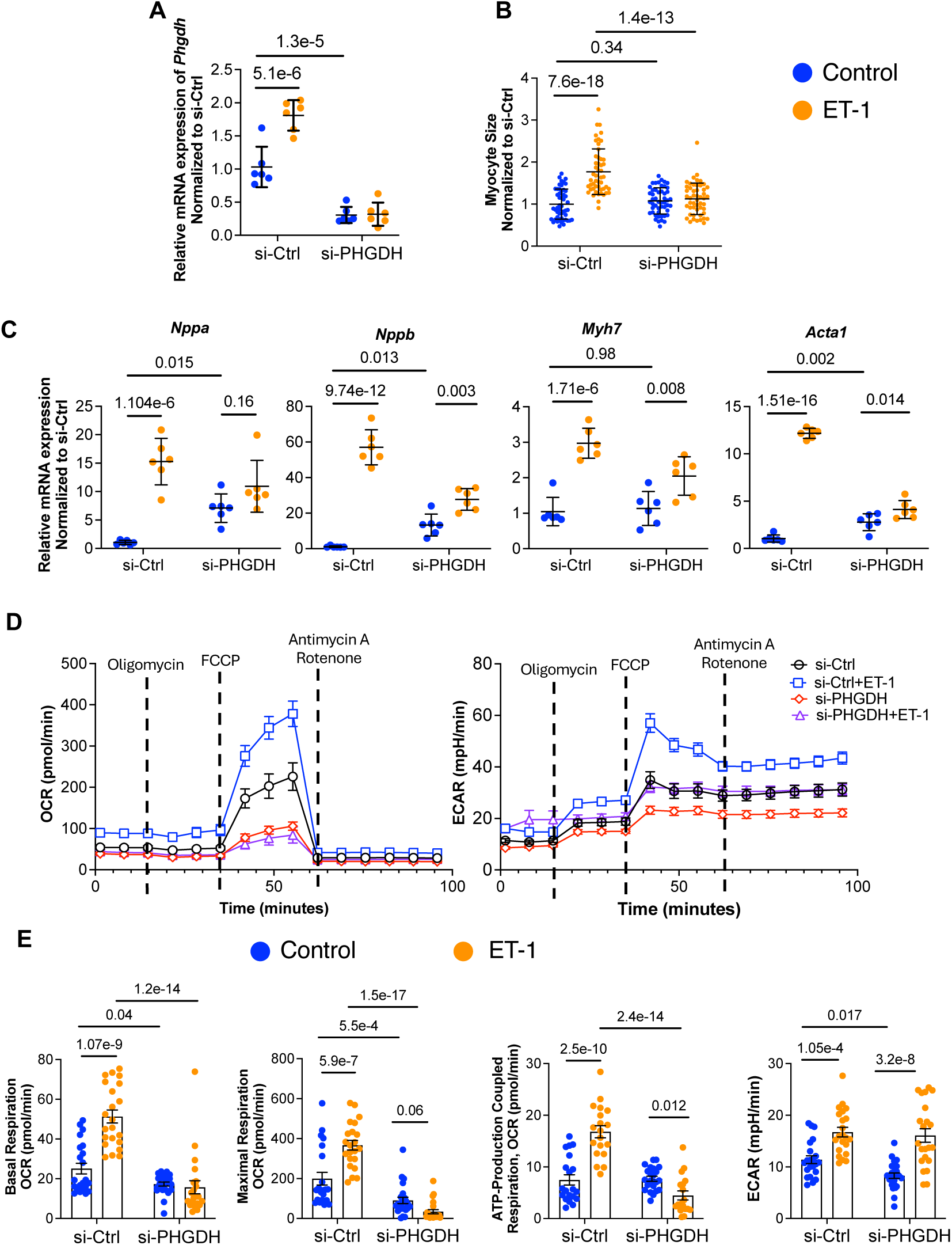
PHGDH suppression blunts ET-1–induced hypertrophic and metabolic remodeling in NRVMs. **A)** *Phgdh* mRNA expression in NRVMs treated with control siRNA (si-Ctrl) or PHGDH siRNA (si-PHGDH), with or without ET-1 stimulation; data normalized to si-Ctrl without ET-1. **B)** Myocyte size in si-Ctrl and si-PHGDH NRVMs treated with or without ET-1; data normalized to si-Ctrl without ET-1. **C)** mRNA expression of hypertrophic markers *Nppa*, *Nppb*, *Myh7*, and *Acta1* in si-Ctrl and si-PHGDH NRVMs treated with or without ET-1; data normalized to si-Ctrl without ET-1. **D)** Seahorse extracellular flux analysis showing oxygen consumption rate (OCR) and extracellular acidification rate (ECAR) in si-Ctrl and si-PHGDH NRVMs treated with or without ET-1, with sequential injection of oligomycin, FCCP, and rotenone/antimycin A. **E)** Summary analysis of basal respiration, maximal respiration, ATP-coupled respiration, and ECAR from the experiment shown in panel D, data shown as mean±SEM.

In control NRVMs, ET-1 also increased basal, maximal, and ATP-linked oxygen consumption rates (OCR) and extracellular acidification rate (ECAR), reflecting enhanced mitochondrial and glycolytic activity due to the stimulation (Figure 4D). Summary data (Figure 4E) shows PHGDH-KD reduced basal and maximal OCR and suppressed ET-1–induced increase in maximal respiratory capacity. ATP-linked respiration also declined in PHGDH-KD cells, consistent with impaired mitochondrial energy production (Figure 4E, lower left). PHGDH suppression also lowered basal ECAR, although ET-1 partially increased ECAR under PHGDH-deficient conditions (Figure 4E, lower right). These results demonstrate that PHGDH is required for cardiomyocyte structural, molecular, and energetic remodeling to hypertrophic stress.

### Cardiomyocyte-targeted PHGDH-KD *in vivo* worsens response to pressure overload

To test whether impaired hypertrophic responses to PHGDH-KD in myocytes translated *in vivo*, we generated cm-specific (*Myh6*-*Cre* x PHGDH^+/flox^) PHGDH^+/−^ KD mice. Genotyping based on the deleted exon 4–5 region confirmed a near 50% reduction in PHGDH^+/-^ (Figure 5A), resulting in a modest but significant (P=0.019) decline in PHGDH protein abundance (Figure 5B). PHGDH^+/-^ and littermate controls (lacking Cre) were subjected to TAC or sham surgery for 4 weeks. At both 1- and 4-week time points in the sham groups, there were no differences between control and PHGDH^+/−^ cardiac structure or function (Supplementary Figure 4A and Figure 5C). However, by as early as 1 week after TAC, systolic adaptation to the stress was abnormal in PHGDH^+/−^ mice, with reduced EF and LV wall thickness, and increased LV systolic volume, consistent with early chamber dilation and systolic dysfunction (Supplementary Figure 5A). This progressed further by 4-weeks after TAC (Figure 5C) with PHGDH^+/-^ hearts developing dilated cardiomyopathy with eccentric rather than concentric hypertrophy. Histology revealed marked fibrosis in the 4-week TAC PHGDH^+/−^ hearts, whereas at baseline, no differences were observed (Figure 5D). 4-week TAC enhanced expression of pathologic gene biomarkers in control and PHGDH^+/−^ mice at generally similar levels, consistent with similar LV mass increase despite differences in geometry (Figure 5E). PHGDH^+/-^ mice also had reduced survival post-TAC (P=0.023) (Figure 5F). There were no significant disparities in group responses related to sex (different symbols in each panel). Since *Myh6* expression increases in the embryo (∼E8) and in turn PHGDH-KD pre-existed TAC, we also tested the effects of PHGDH-KD in a tamoxifen-inducible CM-specific KD model (MerCreMer x with PHGDH^+/flox^), initiating PHGDH knock-down 3-weeks after TAC. This caused similar LV dysfunction/dilation (Supplemental Figure 4B).

**Figure 5:**
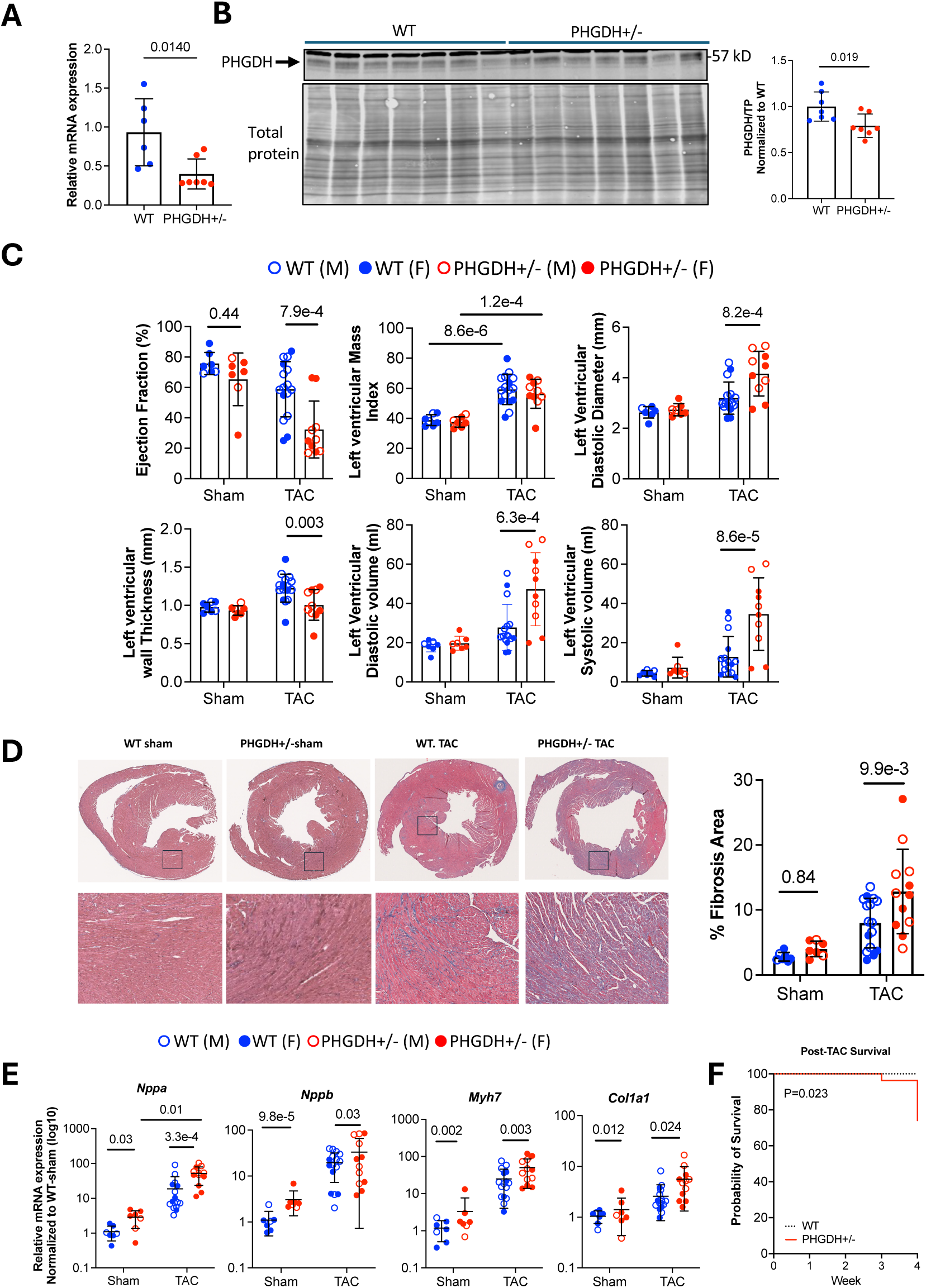
PHGDH haploinsufficiency worsens pressure overload–induced cardiac remodeling and dysfunction. **A)** Validation of reduced myocardial *Phgdh* mRNA expression in PHGDH^+/−^ mice (cardiomyocyte-specific heterozygous PHGDH knockout mice generated using Myh6-Cre) compared with WT controls using primers spanning exons 4–5, the region deleted in this model. **B)** Immunoblot showing myocardial PHGDH and total protein expression in WT and PHGDH^+/−^ mice, with summary densitometry (right) normalized to WT. **C)** Echocardiographic assessment of WT and PHGDH+/− mice following sham or TAC surgery after 4 weeks. **D)** Representative Masson’s trichrome–stained cardiac sections from WT sham, PHGDH^+/−^ sham, WT-TAC, and PHGDH^+/−^-TAC hearts after 4 weeks, with summary quantification of myocardial % fibrosis per total cross-sectional area. **E)** Myocardial mRNA expression of remodeling and fibrosis markers *Nppa*, *Nppb*, *Myh7*, and *Col1a1* in WT and PHGDH+/− mice after sham or TAC surgery after 4 weeks. Data were normalized to WT sham and are presented as log₁₀-transformed values. For panels **C-E,** mouse sex is identified by open circles (male) and closed circles (female). **F)** Kaplan–Meier survival analysis after TAC surgery in WT versus PHGDH^+/−^ mice. There were four deaths in PHGDH^+/−^ mice during weeks 3–4 after TAC: two males and two females.

Lastly, we explored metabolic pathways in PHGDH^+/-^ myocardium before and after TAC to test for translation of the prior cell-based analyses. TAC increased levels of various amino acids in control mice, but they and others were significantly reduced or trended downward in TAC-PHGDH^+/-^ mice (Figure 6A). Similar disparities were observed in members of the folate/methionine/and sulfurination pathways (Figure 6B) with significant reductions in TAC-PHGDH^+/-^ versus TAC-Control in dihydrofolate, 5-methyl tetrahydrofolate, formate, and homocysteine. Reduced glycolytic and TCA intermediates were also observed (Figure 6C) as well as purines (AMP, ADP, and GMP, Figure 6D), several of which had increased after TAC in controls. These data indicate that PHGDH deficiency impairs metabolic adaptation engaged by pressure-overload across a broad range of critical signaling in 1-CM, glycolysis, the TCA cycle, and phosphorylated purines.

**Figure 6:**
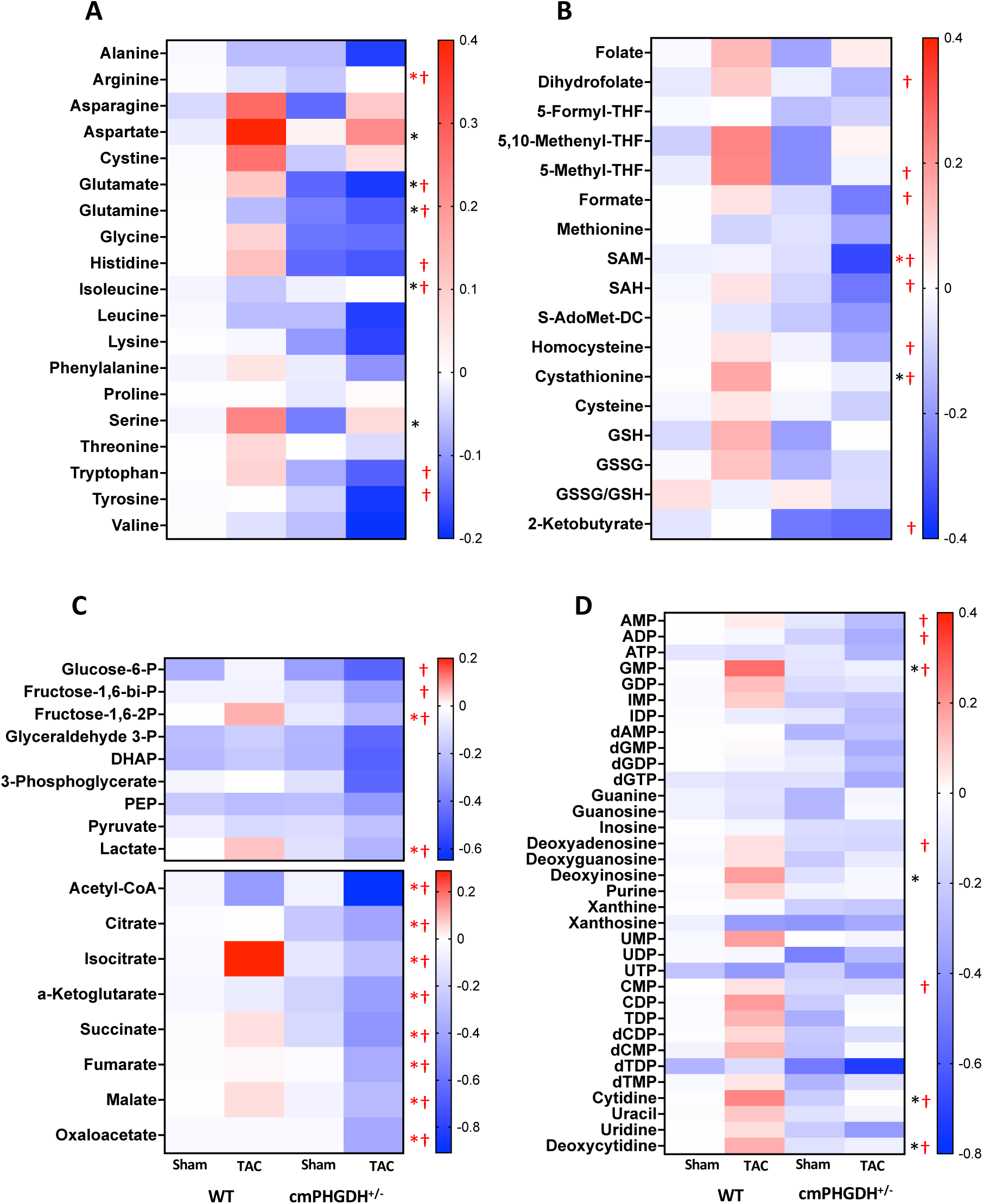
PHGDH haploinsufficiency suppresses metabolic remodeling after pressure overload. **A)** Heat maps of myocardial amino acid abundance, **B)** one-carbon metabolism, serine/glycine metabolism, methionine-cycle, and redox-related metabolites, **C)** glycolytic and tricarboxylic acid cycle metabolites, and **D)** nucleotide and nucleosides and metabolites in WT and PHGDH^+/−^ mice 4 weeks after sham or TAC surgery. Data are normalized to WT sham. Red * indicates PHGDH^+/−^ TAC versus PHGDH^+/−^ sham, red † indicates significant differences between control and TAC for PHGDH^+/-^ mice, and black * indicates the same comparison but in WT mice (*P*<0.1).

## Discussion

This study identifies PHGDH-dependent serine biosynthesis as an important stress-responsive metabolic, anabolic, and energetic pathway required for myocardial adaptation to pathological stress remodeling. Integrating myocardial metabolomics and transcriptomics in humans, and myocyte PHGDH reduction *in vitro* and *in vivo*, we identify endogenous serine biosynthesis as a multi-pronged adaptive metabolic hub linking glycolytic carbon flux with one-carbon metabolism, nucleotide availability, redox buffering, mitochondrial respiration, and anabolic remodeling under cardiac stress. *In vivo*, myocyte PHGDH deficiency impairs coordinated responses to pressure overload, resulting in chamber dilation, systolic dysfunction, fibrosis, and reduced survival.

This study was initially triggered by observations in human HF myocardium, where we found a dissociation between elevated circulating yet depressed myocardial serine. This suggested that systemic serine abundance is insufficient for myocardial serine requirements. Adding to that were findings that key substrates and enzymes for *de novo* serine biosynthesis, 3PG and PHGDH, were substantially reduced. This directed our attention to the SSP and perhaps its biased role in providing one-carbon units for folate-and methionine-cycle activity, and in turn purine and thymidylate synthesis, methylation reactions, and glutathione-dependent redox defense^10,14,30,31^. We found each was compromised by reducing myocyte PHGDH despite exogenous sources, and they were also reduced in metabolomic analysis of human HF myocardium. One feature we did not focus on is the impact of PHGDH regulation of SAM-mediated protein methylation. However, other studies have explored this, reporting serine reduction impacts methylation more via its influences on *de novo* ATP synthesis rather than loss of methyl groups via SAM ^32^. Another study found serine deprivation impairs methylation indirectly by reducing purine-derived ATP needed for methionine to SAM conversion^33^. This suggests the inability to generate sufficient ATP by purinogenesis and mitochondrial alterations is primary.

PHGDH suppression impacted CMs most prominently through three major pathways: nucleotide generation, redox homeostasis, and amino acid formation and associated protein synthesis. The first two appeared the most relevant to CM cytotoxic stress (LDH release), as elevating exogenous amino acids had minimal impact. This indicates PHGDH deficiency does not act solely via depletion of serine, but rather, by selective decline in intracellularly targeted serine and the pathways that depend on this source. This suggests SSP plays a particularly critical role in downstream 1C-metabolism, and in turn purine and redox signaling. This is supported by prior work that found PHGDH inhibition in cancer cell lines impacts nucleotide synthesis via central carbon metabolism, including the pentose phosphate pathway and *de novo* purine and pyrimidine synthesis pathways, with nucleoside supplementation rescuing PHGDH inhibition–induced cellular toxicity ^34^. We too found nucleosides could partially bypass defects from PHGDH-KD, but unlike this prior study, ribose did so as well, and when combined with an antioxidant, the impact was greater in countering LDH release and many 1-CM related pathway defects. Another aspect of CMs that differs from serine handling in cancer cells is the extent to which exogenous serine can overcome suppressed SSP. In cancer, intracellular serine availability is central to purine synthesis and cancer proliferation^15^, but cells readily take up extracellular serine when endogenous synthesis is inadequate or blocked^10,18^. This has stymied the use of PHGDH inhibitors to suppress cancer growth. However, we did not find this in the myocytes, which could explain why human HF exhibits depressed heart serine despite higher plasma serine.

Our metabolomic data indicate that extracellular serine and PHGDH-derived serine play different roles in CM metabolism. Glucose-derived carbon supported central carbon intermediates, amino acid pools, glutathione-related metabolites, and nucleotides. In contrast, exogenous serine contributed more selectively to serine and glycine pools, with a smaller role in glutathione-related metabolites. In cancer, when PHGDH-dependent serine synthesis is constrained by inadequate NAD⁺, extracellular serine uptake bypasses this step to sustain purine biosynthesis ^15^. This does not appear to occur in CMs, as extracellular serine did not substitute for the metabolic defects from SSP suppression.

Both PHGDH-KD and removal of S/G from the media reduced intracellular AA levels, and somewhat surprisingly, this impacted both essential and nonessential AAs. These results show that reduced serine impacts AA balance beyond serine itself, and that reduced serine levels may also affect AA uptake or homeostasis. The influence of PHGDH inhibition on reducing protein synthesis is consistent with prior work in skeletal muscle cells where similar inhibition decreased protein synthesis, myotube size, and myoblast proliferation ^35^. However, the *in vivo* PHGDH^+/-^ model exhibited similarly increased muscle mass after TAC as in controls, so these declines measured over two days are presumably offset more chronically. They could, however, contribute to the lack of cellular hypertrophy over a similar time period in CMs stimulated with ET-1. This aligns with prior studies showing pressure overload activates glycolytic metabolic programs, increasing serine and stimulating YAP during compensatory hypertrophy, and that disruption of these responses promotes dilation and heart failure ^36^.

Coupling of PHGDH-dependent serine biosynthesis and mitochondrial respiratory function, as reported here, is consistent with studies of mitochondrial disease where PHGDH inhibition suppresses oxidative phosphorylation due to compromised 1-carbon metabolism needed for assembly of complex I of the electron transport chain^37^, and generation of sphingolipids needed for inner matrix structures ^38^. This is likely relevant to HF, where mitochondrial dysfunction and impaired stress adaptation are central pathological features ^39^. Complementary data from endothelial cells find PHGDH loss impairs heme synthesis by depleting nucleotides and glutathione, compromising electron transport chain activity, and causing mitochondrial dysfunction and oxidative stress ^40^. In iPSC-derived cardiomyocytes, PHGDH inhibition suppresses complex-I related respiration, increases mitochondrial ROS, and reduces basal and maximal OCR and ATP production, which were partially reversed by N-acetylcysteine, supporting redox dependence ^22^. This too is consistent with the current findings. Collectively, our data support a model in which PHGDH-dependent serine biosynthesis is essential for mitochondrial respiration and antioxidant defense, particularly under high-energy-demand conditions of the heart.

There have been very few *in vivo* studies involving serine manipulation and specifically depletion in the heart, and none to date involving CM-targeted manipulation as in the PHGDH^+/-^ mice in the current study. Our findings support a substantial role of PHGDH and associated SSP in enabling cardiac structural, functional, and metabolic adaptations to cardiac pressure overload. They align with recent gain-of-function studies, finding that PHGDH augmentation is cardioprotective when delivered by AAV9-gene transfer to a model of genetic HFrEF, even without evidence of pre-existing serine or PHGDH deficiency^20^. PHGDH overexpression also promotes cardiomyocyte proliferation that was beneficial in a myocardial infarction model^41^, and analogous benefits were observed by upregulating PSAT1, also in the SSP pathway^21^. The latter was coupled to enhanced nucleotide synthesis, reduced oxidative stress, and limited DNA damage and apoptosis after myocardial injury ^21^. The present study of associated metabolic changes from SSP suppression via PHGDH-KD is, to our knowledge, the first such analysis, and it reveals a major impact on multiple 1C-M related pathways. This supports the centrality of serine homeostasis to myocardial pathophysiology and HFpEF, where it may provide a pleotropic therapeutic node.

Our study has some limitations. Many of the signaling studies were conducted in NRVMs due to their ease of genetic manipulation and subsequent stress testing. However, we also confirmed key features in adult myocytes and, most importantly, in the *in vivo* cm-selective PHGDH^+/-^ model. The metabolic profiling *in vivo* was performed at a single time point, and flux analysis was not performed as was done in myocytes. Additional intensive isotope-tracing studies will be required to measure serine biosynthetic, one-carbon, nucleotide, transsulfuration, and mitochondrial anapleurotic flux in TAC hearts. Rescue studies *in vivo* targeting redox, ribose, or nucleosides, and exogenous serine remain to be performed, though targeting PHGDH itself may indeed be the best therapeutic path. Lastly, several potential downstream mechanisms remain unresolved, including DNA/RNA methylation, histone modifications, mitochondrial respiratory-chain complex activity, mitochondrial dynamics, and lipid remodeling in PHGDH-deficient hearts. These pathways remain important areas for future study.

In conclusion, this study demonstrates that PHGDH-dependent serine biosynthesis is a critical determinant of myocardial stress adaptation. Human heart failure is associated with diminished myocardial serine availability, decreased PHGDH expression, and coordinated remodeling of the serine-glycine one-carbon network. Experimental PHGDH suppression disrupts cardiomyocyte redox balance, nucleotide metabolism, protein synthesis, mitochondrial respiration, and hypertrophic responsiveness. *In vivo* cardiomyocyte-specific PHGDH deficiency impairs the structural, functional, and metabolic adaptations typically activated by pressure overload, resulting in HFrEF-like phenotypes rather than compensatory remodeling. Collectively, these findings identify PHGDH as a central metabolic node that integrates glycolytic carbon flux with one-carbon metabolism, nucleotide supply, redox buffering, and anabolic remodeling, and highlight the serine synthesis pathway as a potential therapeutic target in pathological cardiac remodeling.

## Data availability

All data presented in the manuscript can be made available upon request to the corresponding author.

## Author contributions

Performed experiments and/or data analysis: (MR, MK, NK, TP, SL, DJP, NS, MM). Experimental design: (MR, MK, NK, DAK). Helped with metabolomics studies (LZ, CP, NS). Helped with human studies (KS). Providing data on cardiomyocyte-specific heterozygous *Phgdh* knockout mice generated using the Myh6-CreERT2 model (CH, JS). Wrote and edited manuscript (MR, DAK). Figure preparation (MR, MK, DAK). The authors reviewed and approved the manuscript for publication.

## Acknowledgements

The Gift-of-Life Donor Program, Philadelphia, PA supported procurement of NF-control tissue and HFrEF samples. We thank Drs. Erhe Gao and Nadan Wang in the Small Animal Phenotyping Core at the Johns Hopkins University for performing the animal surgeries and echocardiography. A part of the metabolomics studies were performed by the University of Pennsylvanie Metabolomics Core (RRID:SCR_022381) supported by the Penn Cardiovascular Institute and NCI P30 CA016520 and NIH P30DK050306.

## Abbreviations

PHGDH: phosphoglycerate dehydrogenase
DTT: dithiothreitol
TAC: transverse aortic constriction
HF: heart failure
HFpEF: heart failure with preserved ejection fraction
EF: ejection fraction
HFrEF: heart failure with reduced ejection fraction
LV: left ventricle
3PG: 3-phosphoglycerate
PSAT1: phosphoserine aminotransferase
PSPH: phosphoserine phosphatase
SAM: S-adenosyl methionine
ROS: reactive oxygen species
CM/CMs: cardiomyocyte(s)
NF: non-failing
S/G: serine/glycine
NRVM: neonatal rat ventricular myocyte
KD: knockdown
KO: knockout
NMN: nicotinamide mononucleotide
AAs: amino acids
ET-1: endothelin-1
OCR: oxygen consumption rate
ECAR: extracellular acidification rate
aMHC: alpha myosin heavy chain
1-CM: one-carbon metabolism
SSP: serine biosynthesis pathway (*de novo*)

